# AtlasMap: enabling low-cost, map-style exploration of million-cell single-cell atlases

**DOI:** 10.64898/2026.01.14.699595

**Authors:** Zhou-geng Xu, Hao-Chen Xue, Guihui Qin, Jia-Wei Wang

**Affiliations:** Key Laboratory of Plant Carbon Capture, CAS Center for Excellence in Molecular Plant Sciences (CEMPS), Institute of Plant Physiology and Ecology (SIPPE), Chinese Academy of Sciences (CAS), Shanghai 200032, China; University of Chinese Academy of Sciences, Shanghai 200032, China; Faculty of Health Sciences, University of Macau, Macao SAR, China

## Abstract

Interactive visualization is critical for interpreting single-cell atlases, yet existing web-based tools struggle to handle the growing scale of multimillion-cell datasets, often constrained by browser memory limits and rendering latency. Here, we present AtlasMap, a scalable visualization framework that overcomes these bottlenecks through a multi-resolution, tile-based architecture. Unlike conventional systems that transfer cell-level data to the client, AtlasMap employs an offline preprocessing module to generate quadtree-based spatial aggregations stored in compressed Zarr v3 formats. A high-performance Go backend dynamically renders these summaries as PNG tiles, allowing a lightweight frontend to support fluid pan-and-zoom exploration similar to digital geographic maps. Systematic benchmarking against leading tools (including cellxgene and UCSC Cell Browser) demonstrates that AtlasMap decouples visualization performance from dataset size. While point-based approaches incurred prohibitive client-side memory costs or failed entirely at the 11-million-cell scale, AtlasMap maintained sub-second startup latency and a negligible browser footprint (<5 MB). By shifting computational pressure from the browser to an optimized server-side pipeline, AtlasMap enables accessible, high-fidelity exploration of ultra-scale single-cell datasets on standard hardware.

## Introduction

Recent advances in single-cell RNA sequencing (scRNA-seq) have unveiled cellular heterogeneity at unprecedented resolution, driving the construction of massive cellular atlases across diverse tissues, disease contexts, and species(Jovic et al. 2022). As these initiatives expand in both breadth and depth, datasets increasingly encompass millions of cells, thousands of genes, and rich metadata layers ranging from donor information to multi-level annotations(Xu et al. 2023). While these resources are pivotal for biological discovery and reference mapping, their sheer scale and complexity have significantly raised the barrier to data dissemination and reuse, rendering traditional “download-then-analyze” workflows increasingly prohibitive.

To democratize access to these resources, a diverse ecosystem of scRNA-seq analysis and visualization tools has emerged. Existing solutions generally fall into two categories: integrated platforms that couple preprocessing, analysis, and visualization within a unified environment(Long et al. 2025; Weinreb et al. 2018; Pan et al. 2025; Bya et al. 2025); and specialized browser-based systems focused on the interactive exploration and sharing of precomputed results (e.g., cellxgene(CZI Cell Science Program et al. 2025) and UCSC Cell Browser(Speir et al. 2021)). These browser-centric tools play a crucial role in lowering the technical threshold for non-computational users, facilitating rapid inspection of embedding spaces, gene expression patterns, and metadata distributions(Xue et al. 2025). Concurrently, the growing necessity for consortium-level and cross-species atlas integration places higher demands on the deployability and scalability of these visualization systems.

Among browser-based tools, architectural designs differ fundamentally in their data organization and serving paradigms. One class of tools operates by directly loading and serving cell-level objects during the online phase, prioritizing ease of use and streamlined workflows. The other class relies on offline preprocessing to generate precomputed artifacts—such as aggregated summaries or multi-resolution representations—which are then served on-demand. While both paradigms offer acceptable performance for smaller datasets, interactive visualization becomes a critical bottleneck at the atlas scale. When the browser is tasked with rendering millions of scatter points and supporting dynamic re-coloring, filtering, or zooming, the costs of client-side rendering and data transmission scale linearly with cell count. This results in high initial latency, prohibitive memory spikes, and degraded responsiveness, particularly on resource-constrained devices or limited networks.

Here, we introduce AtlasMap, a scalable framework for the interactive visualization of large-scale scRNA-seq embeddings designed to alleviate frontend rendering pressure and bandwidth limitations while supporting unified multi-project deployment. Beyond presenting the system architecture, we establish a systematic evaluation framework based on these execution paradigms to benchmark representative visualization tools. By decoupling evaluation into preprocessing and online phases, we characterize how computational resource peaks and time costs shift between server and client at standard scales (1K–1M cells). Furthermore, we stress-test these systems in an ultra-scale scenario (11M cells) to identify the failure modes and critical constraints that define the boundaries of current visualization feasibility.

## Result

### Architecture and execution models determine where computational pressure emerges

To contextualize differences observed in the subsequent benchmarks, we first compared workflow-level design choices across five single-cell visualization systems (Table 1). We focused on factors that determine where computational and memory pressure emerges along the end-to-end pipeline: whether offline preprocessing is required, what the online service is responsible for, the granularity of data delivered to the client, and whether an explicit level-of-detail (LOD) mechanism constrains interactive retrieval across zoom levels. This comparison reveals two dominant execution paradigms:direct-loading systems that operate primarily on cell-level objects during online serving (cellxgene and cirrocumulus), and offline-preprocessed systems that shift substantial cost to an explicit preprocessing stage and serve derived artifacts during browsing (AtlasMap, UCSC Cell Browser, and ShinyCell2).

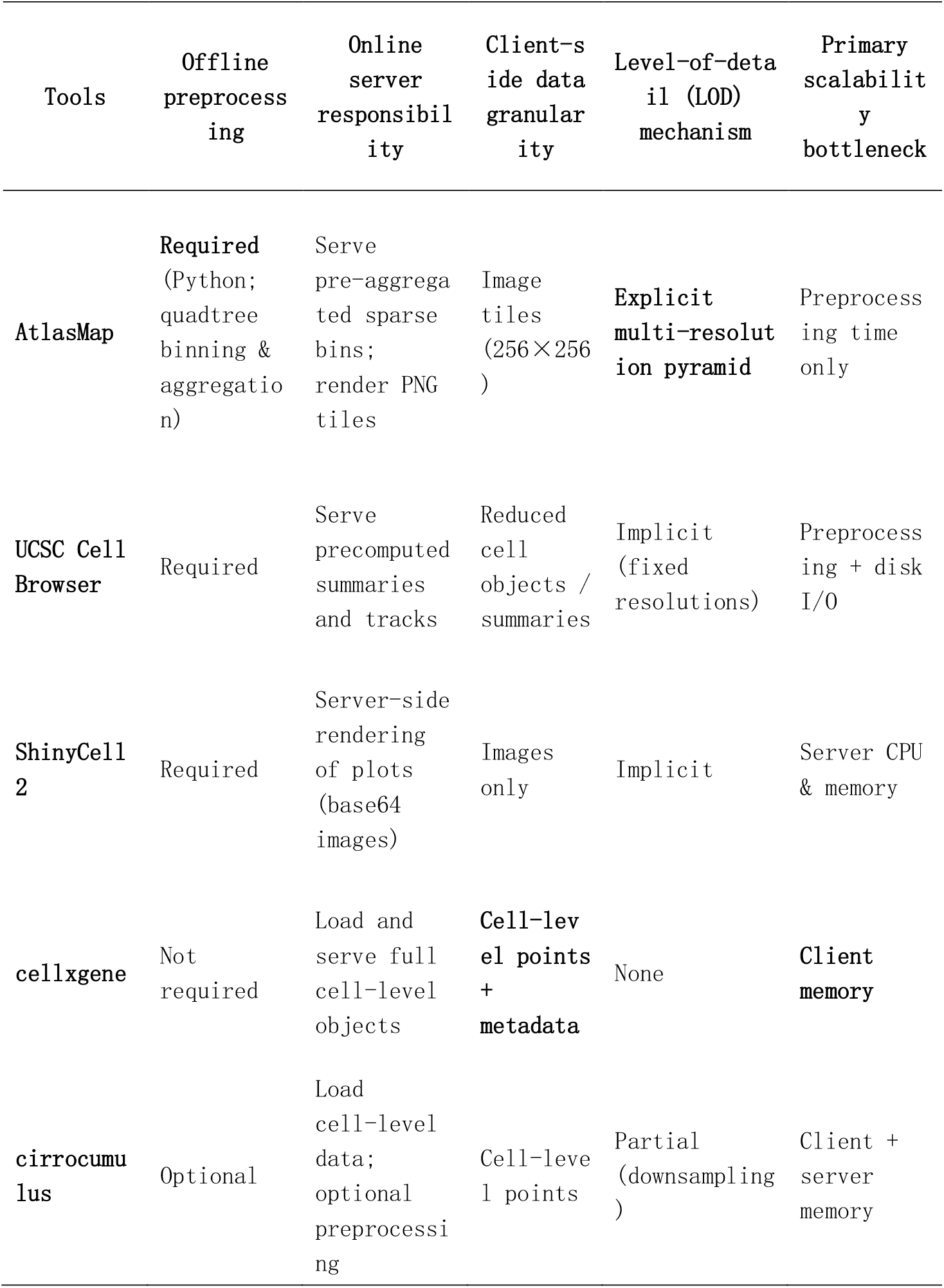
Table 1.

**Table1**
| Tools | Offline preprocessing | Online server responsibility | Client-side data granularity | Level-of-detail (LOD) mechanism | Primary scalability bottleneck |
| --- | --- | --- | --- | --- | --- |
| AtlasMap | Required (Python; quadtree binning & aggregation) | Serve pre-aggregated sparse bins; render PNG tiles | Image tiles (256×256) | Explicit multi-resolution pyramid | Preprocessing time only |
| UCSC Cell Browser | Required | Serve precomputed summaries and tracks | Reduced cell objects / summaries | Implicit (fixed resolutions) | Preprocessing + disk I/O |
| ShinyCell 2 | Required | Server-side rendering of plots (base64 images) | Images only | Implicit | Server CPU & memory |
| cellxgene | Not required | Load and serve full cell-level objects | Cell-level points + metadata | None | Client memory |
| cirrocumulus | Optional | Load cell-level data; optional preprocessing | Cell-level points | Partial (downsampling) | Client + server memory |

Among direct-loading tools,cellxgene does not require preprocessing and is designed to load and serve cell-level data during the online phase, with the browser largely interacting with point-level coordinates and associated metadata. Cirrocumulus shares a similar online-first structure but supports optional preprocessing, providing a configurable path between immediate direct loading and an upfront preparation step. In practice, this flexibility implies that the dominant resource bottleneck (startup, steady-state serving, or interactive browsing) can shift with dataset scale and deployment configuration, even within the same system.

In contrast, offline-preprocessed tools explicitly adopt a segmented preprocess-serve-browser pipeline and rely on precomputed outputs during online interaction (Table 1). UCSC Cell Browser and ShinyCell2 both require preprocessing and distribute browser-ready assets at runtime; notably, ShinyCell2 primarily delivers server-rendered plot images (e.g., base64-encoded) rather than client-side rendered point layers, shifting interactive rendering work toward the backend. AtlasMap also requires preprocessing, but differs in the unit of interaction delivered to the client: instead of exposing cell-level coordinates as the primary payload, it serves tile-based, multi-resolution summaries and is the only tool in this comparison that implements an explicit LOD mechanism. These workflow distinctions—cell-level payloads, server-rendered images, and multi-resolution summaries—provide a natural explanation for why pressure concentrates at different stages across tools. Accordingly, the subsequent analysis quantifies performance along three axes aligned with these execution paradigms: server-side pressure, browser-side pressure, and ultra-scale feasibility.

### Preparation-phase scaling shifts computational cost between runtime and preprocessing

To compare the computational pressure incurred before interactive exploration is possible, we report time and peak memory during a unified dataset preparation stage (Methods). For offline-preprocessed workflows (AtlasMap, ShinyCell2, and UCSC Cell Browser), this stage corresponds to the explicit preprocessing/compilation pipeline. For direct-loading workflows (cellxgene and cirrocumulus), we treat the initial web-loading phase as an implicit preparation step, reflecting computation and data materialization required before the dataset becomes explorable in the browser.

Across 1K–1M cells, preparation-stage peak memory and elapsed time diverged markedly by workflow (Fig. 1). At the 1M scale, direct-loading tools completed the preparation stage in 61.84 s (cellxgene) and 32.49 s (cirrocumulus), with peak RSS of 24.4 GB and 13.7 GB, respectively. In contrast, offline-preprocessed workflows shifted the dominant cost into preprocessing: AtlasMap completed preprocessing in 418.13 s with 58.2 GB peak RSS, UCSC Cell Browser required 1452.10 s with 47.7 GB peak RSS, and ShinyCell2 required 3052.11 s with 116.5 GB peak RSS. These results indicate that workflow architecture primarily determines when the dominant memory/time cost is paid: direct-loading tools incur a smaller but online preparation cost, whereas offline pipelines concentrate larger peaks into a one-time preprocessing step.

**Figure 1.**
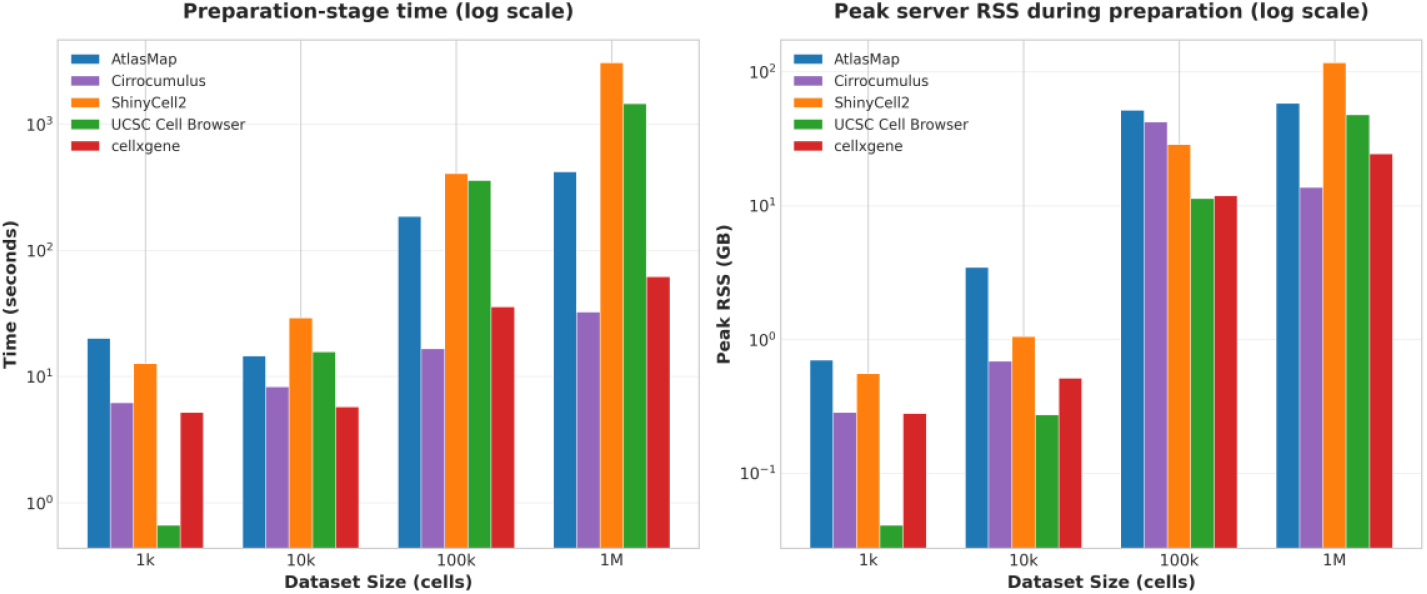
Preparation-stage time and peak server memory across conventional dataset scales (1K-1M) Left, wall-clock time required to prepare a dataset before interactive exploration. Right, peak process-tree resident set size (RSS) measured during the same preparation interval (reported in GB; log-scaled y-axes). For cross-tool comparability, we define a unified dataset preparation stage as the minimal computation required before a dataset becomes explorable in the browser: for offline-preprocessed workflows (AtlasMap, ShinyCell2, and UCSC Cell Browser), this corresponds to the explicit preprocessing/compilation pipeline; for direct-loading workflows (cellxgene and cirrocumulus), which do not have an offline preprocessing step, we treat the initial web-loading interval (service launch through first successful page load) as an implicit preparation step. For UCSC Cell Browser, values reflect the dataset compilation step used to generate deployable browser artifacts.

### Web-phase scaling reveals high client requirements for point-based loading

We next quantified web-phase performance after deployment using three complementary metrics: startup latency, peak server RSS, and browser JavaScript heap (Fig. 2). AtlasMap exhibited nearly size-invariant behavior from 1K to 1M cells, with 0.009–0.014 s startup latency, 126–138 MB peak server RSS, and 4.89–4.96 MB client heap. Among offline-preprocessed baselines, ShinyCell2 maintained similarly small client heaps (5.54–6.08 MB) but incurred higher startup latency (∼4.18–4.43 s) and a larger server footprint (402–1,074 MB). In contrast, point-based direct-loading workflows increased both server and client pressure with dataset size: at 1M cells, cellxgene reached 58.67 s startup latency with 25.23 GB peak server RSS and 462.65 MB client heap, while cirrocumulus preserved short startup times (∼1.42 s) but still required 17.66 GB peak server RSS and 152.06 MB client heap. Notably, UCSC Cell Browser maintained a minimal server footprint (∼55 MB) yet imposed a high client heap that increased up to 411.42 MB at 1M cells, indicating a workflow where substantial state is materialized in the browser rather than the server.

**Figure 2.**
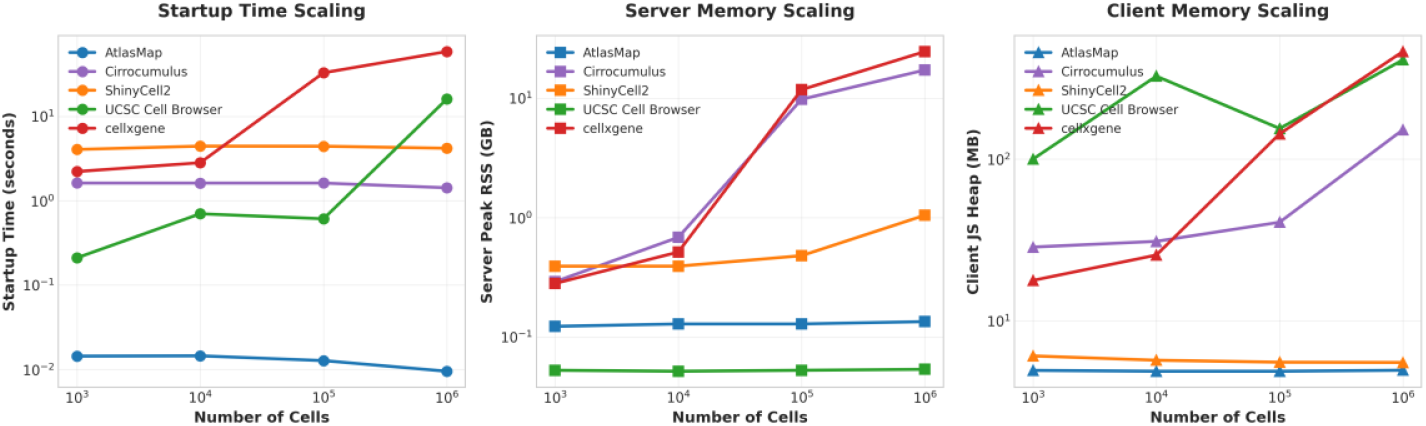
Web-phase scaling of latency and memory across conventional dataset sizes (1K–1M). The deployed web instances was evaluated by measuring (i) startup latency (startup_s, wall-clock time from navigation to a ready page state), (ii) peak server memory (peak process-tree RSS during the same interval), and (iii) client memory (JavaScript heap after page load). Across all scales, AtlasMap maintained near-constant server RSS (∼0.13 GB) and client heap (∼5 MB) with sub-second startup latencies. In contrast, point-based direct-loading tools showed pronounced growth in server and/or client footprints, reaching tens of GB server RSS and hundreds of MB client heap at 1M cells.

In typical exploratory sessions, users often prioritize inspecting expression patterns for a small number of genes (e.g., marker genes) over rendering every cell point at maximal fidelity. Under this usage model, the key usability bottleneck becomes how quickly the interface reaches an explorable embedding and supports the first expression overlay—a behavior well approximated by our startup latency measurement. The observed scaling therefore implies a practical constraint: workflows that require hundreds of MB of client heap at the million-cell scale (e.g., cellxgene and UCSC in this benchmark) place substantially higher demands on end-user browsers and may exhibit degraded responsiveness, whereas tile-summary-based clients (AtlasMap; and ShinyCell2 in this setting) maintain low and stable browser memory footprints, enabling faster time-to-first-view.

### Ultra-scale benchmarking exposes feasibility limits and failure modes

We next stress-tested all workflows on the 11M-cell dataset to determine practical feasibility limits and failure modes. At this scale, tools diverged sharply in where they failed—during preprocessing, server-side stabilization, or client-side rendering.

ShinyCell2 could not be evaluated because conversion to a Seurat object fails during preprocessing: the R sparse-matrix backend imposes a 2^31^−1 (∼2.15×10^9^) non-zero entry limit, which is exceeded at 11M cells. Cirrocumulus required an extended preprocessing/startup phase of 3 h 28 min and reached a peak RSS of 1,577.84 GB; after completion, server memory remained highly elevated and fluctuated between 458.16–930.42 GB. Once accessible, page refreshes completed in ∼25 s and browser JS heap stabilized at ∼968 MB. cellxgene stabilized server memory after ∼12 min at ∼228.53 GB but exhibited rapid client-side growth: JS heap reached ∼1.44 GB within ∼30 s and continued increasing until the browser became unresponsive or crashed. The UCSC Cell Browser exhibited significant computational demands. The compilation process (cbBuild) peaked at 294 GB RSS and typically required approximately 6 hours to complete. Furthermore, reloading the web interface took roughly 7.8 minutes, necessitating 466 MB of network data transfer and occupying 1.54 GB of the frontend JavaScript heap memory

In contrast, AtlasMap completed preprocessing in 900 s with a peak RSS of 160 GB and remained responsive at deployment: the web page opened in <1 s and client JS heap stayed <5 MB. Collectively, these results show that ultra-scale feasibility is jointly constrained by preprocessing cost, server memory requirements, and-critically for point-based workflows-client-side stability under large-scale rendering.

## Discussion

We conducted an end-to-end benchmark of five interactive single-cell visualization systems, focusing on the pipeline comprising preparation, online serving, and browser-side rendering. Our primary objective was to investigate where computation and state are materialized and how this placement impacts resource utilization and scalability. Our results indicate that system performance cannot be captured by a single metric. Instead, the tools fall into two distinct architectural paradigms: (1) Direct Loading with Point-Level Rendering (e.g., cellxgene, cirrocumulus), which concentrates costs during the online phase by maintaining cell-level objects on both the server and client; and (2) Offline Preprocessing with Summary/Slice Browsing (e.g., AtlasMap, UCSC Cell Browser, ShinyCell2), which shifts computational burdens to a one-time preparation step, serving primarily derived artifacts—such as image tiles, pre-computed structures, or summaries—during browsing(Keller et al. 2025; Schaefer et al. 2025). This dichotomy consistently explains the divergent behaviors observed in server memory, client memory, and usability at ultra-large scales.

At standard scales (1K–1M cells), client-side memory emerges as a critical differentiator of usability. Point-level workflows compel the browser to materialize and retain substantial client-side state. For instance, at 1M cells, the JavaScript (JS) heap size reached hundreds of megabytes (cellxgene: ∼462.65 MB; UCSC Cell Browser: ∼411.42 MB; cirrocumulus: ∼152.06 MB). In contrast, AtlasMap, utilizing a tile/summary approach, maintained a nearly constant client heap (∼5 MB) across the 1K–1M range. This constraint becomes definitive at ultra-large scales (11M cells). Although point-level systems maintained feasible memory footprints on the server (e.g., cellxgene: ∼228.53 GB), the browser failed due to unbounded heap growth (reaching ∼1.44 GB within 30 seconds before crashing). Similarly, UCSC Cell Browser, despite utilizing offline preprocessing, became unstable due to the bottleneck of client-side point-level rendering. These observations underscore that at the multi-million cell level, “server feasibility” does not equate to “user availability,” with browser stability acting as the decisive bottleneck for interaction.

From a user-centric perspective, exploratory analysis often focuses on examining the expression of a few marker genes. Consequently, metrics such as “time-to-first-browse” and “time-to-first-expression” are more representative of usability than the immediate rendering of all cells at maximum fidelity. Consistent with this, AtlasMap achieved sub-second page loads at 11M cells with minimal client overhead, demonstrating the scalability benefits of summary-first and Level-of-Detail (LOD) strategies. Conversely, point-level approaches (including offline tools that retain point rendering) offer advantages in cell-by-cell selection semantics and immediate fidelity; however, their client-side resource demands grow linearly with dataset size, limiting feasibility on ultra-large datasets. Furthermore, decoupling the frontend and backend concentrates heavy computational loads into amortizable one-time preprocessing steps (e.g., AtlasMap peaked at ∼160 GB during 11M preprocessing). This significantly reduces the resident resources required for online serving and associated cloud costs, thereby lowering deployment barriers for small laboratories and collaborative scenarios. It is important to note that ultra-large-scale analysis is also constrained by ecosystem and data representation limits (e.g., ShinyCell2 failed at 11M due to the 231−12^{31}-1231−1 non-zero element limit in R sparse matrices). Additionally, while this benchmark primarily covered embedding visualization and expression overlays, future work should extend to richer interactive tasks and systematically incorporate user-centric metrics.

## Methods

### Software architecture

The system implements a tile-based architecture for scalable interactive single-cell visualization. An offline preprocessing module (Python) ingests 2D embeddings (e.g., UMAP) and performs quadtree-based spatial binning with sparse aggregation to generate multi-resolution summaries. The processed outputs are stored as compressed Zarr v3 artifacts organized as a multi-resolution pyramid of pre-aggregated bins. A high-performance backend service (Go) reads the required sparse blocks on demand, caches frequently accessed results, and renders 256×256 PNG tiles for client requests. A TypeScript frontend built on Leaflet displays the tiles as interactive map layers, enabling smooth pan/zoom exploration across dataset scales.

### Data collection

Single-cell RNA-seq datasets were obtained from the cellxgene data portal. We selected publicly available datasets spanning different tissues, organisms, assay types, and scales (from ∼10^3^ to >10^7^ cells), including pancreas, blood, brain, and embryonic development samples from *Danio rerio, Homo sapiens*, and *Mus musculus*.

### Software configuration

To ensure reproducibility and avoid dependency conflicts, each visualization tool was installed in an isolated conda environment using mamba with the conda-forge channel under strict priority. We evaluated four commonly used single-cell visualization frameworks: cellxgene(CZI Cell Science Program et al. 2025), UCSC Cell Browser(Speir et al. 2021), ShinyCell2(Ouyang et al. 2021), and cirrocumulus(Li et al. 2020).

For cellxgene and cirrocumulus, Python 3.11 environments were created and the latest stable releases were installed via pip; datasets were launched directly from .h5ad files using the respective command-line interfaces. UCSC Cell Browser was installed in a separate Python environment, with .h5ad files imported using cbImportScanpy followed by dataset compilation using cbBuild. ShinyCell2 was installed in an R environment with binary dependencies provided by conda-forge; .h5ad files were converted to Seurat objects and preprocessed using schard and ShinyCell2 utilities to generate standalone Shiny applications.

All tools were tested using the same minimal dataset (1K cells) to validate successful installation and execution prior to large-scale benchmarking, and complete environment specifications were exported to YAML files for reproducibility.

### Runtime memory profiling in Python

To quantify runtime memory overhead, each generated .h5ad file was loaded with Scanpy (Wolf et al. 2018)(sc.read_h5ad) and the process resident set size (RSS) was recorded using psutil immediately before and after loading (with garbage collection). For each dataset, we reported file size on disk, cell/feature dimensions, load time, and the RSS increase attributable to loading (MB), together with the memory-to-file-size ratio.

### Runtime memory profiling in R

To quantify memory overhead for Seurat-based workflows, we performed an R-side benchmark analogous to the Scanpy .h5ad loading test. For each dataset, the .h5ad file was converted to a Seurat object using schard::h5ad2seurat and serialized to a compressed .qs file (qs::qsave) to enable repeatable load-time measurements. Due to a known limitation of the Matrix sparse format used by Seurat (a 2^31−1 cap on the number of non-zero entries), the 11M cell dataset could not be converted into a Seurat object and was therefore excluded from the Seurat benchmark. We measured resident set size (RSS) using the ps package immediately before and after loading the .qs file (qs::qread), with garbage collection invoked to reduce transient allocations. For each dataset, we reported on-disk file sizes (.h5ad and .qs), .qs load time, RSS increase attributable to loading, and basic dataset dimensions (cells and genes).

### Server-side memory profiling during preprocessing

To quantify server memory consumption during dataset preprocessing, we instrumented each preprocessing pipeline with a lightweight RSS profiler that samples the resident set size of the full process tree (parent process plus all descendants). For each preprocessing run, the profiler launched the preprocessing command in its own process group, sampled per-process resident memory via the Linux procfs statm interface at a fixed interval, and summed RSS across all descendant processes to report peak memory usage.

### Browser-side performance evaluation

Client-side metrics were collected using an automated headless Chrome workflow implemented in Puppeteer. For each tool instance, Puppeteer navigated to the local deployment URL and recorded standard Navigation Timing entries from the Performance API (including timing components related to request/response and page readiness). Where supported by the browser, JavaScript heap usage was additionally captured via performance.memory (used/total heap and heap limit). To approximate interactive usage, the script optionally performed standardized wheel-based zoom in/out events on a target element (default: canvas) and logged whether the interaction target was detected and successfully exercised. All metrics were exported as structured JSON for downstream aggregation and comparison across tools and dataset sizes.

### Benchmark metrics

Tools were benchmarked under a unified framework consisting of two distinct phases: preprocessing and runtime.

For offline-preprocessed tools (AtlasMap, ShinyCell2, UCSC Cell Browser), the preprocessing phase comprised the execution of data conversion commands (e.g., cbBuild), measuring execution time and peak server RSS. For direct-loading tools (cellxgene, cirrocumulus), we redefined the preprocessing phase to include the interval from service launch to the completion of the initial browser page load; metrics tracked were time-to-load, peak server RSS, and initial browser JS heap.

The runtime phase evaluated performance after the web service was fully established (or after the initial load for direct tools). During this phase, we measured the stable server-side RSS and the client-side JS heap usage (via Puppeteer) to assess the memory burden during active interaction. This alignment allows for a direct comparison of initialization overheads versus steady-state resource consumption across all tools.

## Acknowledgments

We thank Shipeng Guo and Jianing Gao for their valuable suggestions and discussions.

## Author contributions

Z-G.X. conceived the study, developed the core algorithms, and drafted the initial manuscript. G.Q. collected and organized the single-cell datasets used for benchmarking. H-C.X. conducted the comprehensive software usability evaluation and performance testing. J-W.W. provided critical feedback, reviewed the manuscript, and supervised the project.

## Funding

This work was supported by the Postdoctoral Fellowship Program of CPSF (GZB20230757), and the Special Research Assistant of CAS.

## Data and code availability

The source code for AtlasMap is publicly available on GitHub at https://github.com/xuzhougeng/atlasmap-sc. The benchmarking and evaluation scripts used in this study are also available on GitHub at https://github.com/xuzhougeng/atlasmap-benchmark.

## Supplementary data

## Notes

### Competing Interest Statement

The authors have declared no competing interest.

## References

Bya, Phi, Duy Tran, Khoi Nguyen, Sorin Draghici, and Tin Nguyen. 2025. “CytoAnalyst Web Platform Facilitates Comprehensive Single Cell RNA Sequencing Analysis.” Scientific Reports 15 (1): 28736. 10.1038/s41598-025-14398-x.

CZI Cell Science Program, Shibla Abdulla, Brian Aevermann, et al. 2025. “CZ CELLxGENE Discover: A Single-Cell Data Platform for Scalable Exploration, Analysis and Modeling of Aggregated Data.” Nucleic Acids Research 53 (D1): D886–900. 10.1093/nar/gkae1142.

Jovic, Dragomirka, Xue Liang, Hua Zeng, Lin Lin, Fengping Xu, and Yonglun Luo. 2022. “Single-Cell RNA Sequencing Technologies and Applications: A Brief Overview.” Clinical and Translational Medicine 12 (3): e694. 10.1002/ctm2.694.

Keller, Mark S., Ilan Gold, Chuck McCallum, Trevor Manz, Peter V. Kharchenko, and Nils Gehlenborg. 2025. “Vitessce: Integrative Visualization of Multimodal and Spatially Resolved Single-Cell Data.” Nature Methods 22 (1): 63–67. 10.1038/s41592-024-02436-x.

Li, Bo, Joshua Gould, Yiming Yang, et al. 2020. “Cumulus Provides Cloud-Based Data Analysis for Large-Scale Single-Cell and Single-Nucleus RNA-Seq.” Nature Methods 17 (8): 793–98. 10.1038/s41592-020-0905-x.

Long, Renwen, Tina Suoangbaji, Daniel Wai-Hung Ho, Renwen Long, Tina Suoangbaji, and Daniel Wai-Hung Ho. 2025. “OmniCellX: A Versatile and Comprehensive Browser-Based Tool for Single-Cell RNA Sequencing Analysis.” Biology 14 (10). 10.3390/biology14101437.

Ouyang, John F, Uma S Kamaraj, Elaine Y Cao, and Owen J L Rackham. 2021. “ShinyCell: Simple and Sharable Visualization of Single-Cell Gene Expression Data.” Bioinformatics 37 (19): 3374–76. 10.1093/bioinformatics/btab209.

Pan, Rufei, Chao Li, Feifei Shi, et al. 2025. “BestopCloud: An Integrated One-Stop Solution for Single-Cell RNA Sequencing Data Analysis.” BMC Genomics 26 (1): 905. 10.1186/s12864-025-12072-0.

Schaefer, Moritz, Peter Peneder, Daniel Malzl, et al. 2025. “Multimodal Learning Enables Chat-Based Exploration of Single-Cell Data.” Nature Biotechnology, November 11, 1–11. 10.1038/s41587-025-02857-9.

Speir, Matthew L, Aparna Bhaduri, Nikolay S Markov, et al. 2021. “UCSC Cell Browser: Visualize Your Single-Cell Data.” Bioinformatics 37 (23): 4578–80. 10.1093/bioinformatics/btab503.

Weinreb, Caleb, Samuel Wolock, and Allon M Klein. 2018. “SPRING: A Kinetic Interface for Visualizing High Dimensional Single-Cell Expression Data.” Bioinformatics 34 (7): 1246–48. 10.1093/bioinformatics/btx792.

Wolf, F. Alexander, Philipp Angerer, and Fabian J. Theis. 2018. “SCANPY: Large-Scale Single-Cell Gene Expression Data Analysis.” Genome Biology 19 (1): 15. 4801. 10.1186/s13059-017-1382-0.

Xu, Chuan, Martin Prete, Simone Webb, et al. 2023. “Automatic Cell-Type Harmonization and Integration across Human Cell Atlas Datasets.” Cell 186 (26): 5876–5891.e20. 25. 10.1016/j.cell.2023.11.026.

Xue, Hao-Chen, Zhou-Geng Xu, Yu-Jie Liu, et al. 2025. “A Unified Cell Atlas of Vascular Plants Reveals Cell-Type Foundational Genes and Accelerates Gene Discovery.” Cell 188 (22): 6370–6390.e29. 10.1016/j.cell.2025.07.036.

